# The adaptive role of cell death in yeast communities stressed with macrolide antifungals

**DOI:** 10.1101/2021.08.27.457932

**Authors:** N. Kireeva, S.S. Sokolov, E.A. Smirnova, K.V. Galkina, F.F. Severin, D.A. Knorre

## Abstract

Microorganisms cooperate with each other to protect themselves from environmental stressors. An extreme case of such cooperation is regulated cell death for the benefit of other cells. Dying cells can provide surviving cells with nutrients or induce their stress-response by transmitting an alarm signal; however, the role of dead cells in microbial communities is unclear. Here we searched for types of stressors the protection from which can be achieved by death of a subpopulation of cells. Thus, we compared the survival of *Saccharomyces cerevisiae* cells upon exposure to various stressors in the presence of additionally supplemented living versus dead cells. We found that dead cells contribute to yeast community resistance against macrolide antifungals (e.g. amphotericin B [AmB] and filipin) to a greater extent than living cells. Dead yeast cells absorbed more macrolide filipin than control cells because they exposed intracellular sterol-rich membranes. We also showed that, upon the addition of lethal concentrations of AmB, supplementation with AmB-sensitive cells but not with AmB-resistant cells enabled the survival of wild-type cells. Together, our data suggests that cell-to-cell heterogeneity in sensitivity to AmB can be an adaptive mechanism helping yeast communities to resist macrolides, which are naturally occurring antifungal agents.

**Importance:** Eukaryotic microorganisms harbour elements of programmed cell death (PCD) mechanisms that are homologous to the PCD of multicellular metazoa. However, it is still debated whether microbial PCD has an adaptive role or the processes of cell death are an aimless operation in self-regulating molecular mechanisms. Here, we demonstrated that dying yeast cells provide an instant benefit for their community by absorbing macrolides, which are bacteria-derived antifungals. Our results illustrate the principle that the death of a microorganism can contribute to the survival of its kin and suggest that early plasma membrane permeabilization improves community-level protection. The latter makes a striking contrast to the manifestations of apoptosis in higher eukaryotes, the process by which plasma membranes maintain integrity.

## Introduction

Microorganisms compete with each other for resources, and the success of this competition depends on their ability to resist toxic compounds produced by the contenders. The resistance can be induced by multiple mechanisms. For example, cells can prevent uptake of toxic inhibitor molecules (1), actively efflux toxic compounds with plasma membrane transporters (2) or metabolise toxic molecules (3). Meanwhile, different mechanisms can cause opposite effects on surrounding cells. On one hand, drug efflux reduces the concentration of the xenobiotic in the cytoplasm, but this does not help neighbouring cells to withstand the stress. On the other hand, if a microorganism decomposes a xenobiotic, it produces a ‘common good’ by increasing the chances of surrounding cells to survive (see (4) for review). To better exploit these cooperative mechanisms, microbial cells form multicellular aggregates or biofilms. For example, bacterial cells treated with sub-lethal concentrations of antimicrobial peptides induce cell aggregation (5). In yeast, flocculation increases cellular tolerance to macrolide antifungal amphotericin B (AmB) and hydrogen peroxide, despite that functional flocculin allele *FLO11* decreases individual cell fitness (6). Moreover, some yeast strains form colonies with different cell layers, and cells in the exterior layer show increased resistance to environmental stressors (7).

An extreme level of microbial cooperation is ‘altruistic’ death, a process by which cells die to provide their neighbouring cells with nutritional and environmental conditions that support their survival (8–10). The death of some cells in microbial suspension or biofilm can provide an advantage to surviving cells in different ways. For example, it has been shown that cell death in *Escherichia coli* mediated by the *mazF* module of *maxEF* toxin/antitoxin system can prevent the spread of the phages across the bacterial population (11). Moreover, individual *E. coli* cells have different bacterial toxin production rates, so some cells produce more toxins than the others, autolyse and release the toxin to the medium. This toxin inhibits other bacterial strains that lack this toxin/antitoxin system (12). This strategy is considered to be an adaptive manifestation of microbial cell lysis. Furthermore, inviable cells still increase the fitness of their kin by providing them with nutrients (13, 14) or transmitting an alarm signal that induces adaptation to stress in surviving cells (15, 16). In the case of pathogens, dead cells can modulate the host immune response, thereby preparing a niche for further invasion (17). Finally, dead cells can absorb toxic compounds while allowing surviving cells to continue proliferation (18). Interestingly, it has been shown that dead algae cells absorb pollutants better than living cells (19). Together, this indicates that, although dead cells cannot propagate their genes to offspring, the biochemical processes in their remnants can have a significant effect on the survival of surrounding cells.

Meanwhile, some environmental stressors or xenobiotics induce yeast death that can be prevented by the inhibition of regulatory cascades (20–24). For example, the deletion of metacaspase or endonuclease G genes, which are homologues of mammalian apoptosis transducers, prevents yeast death induced by oxidative stress (22, 25). However, whether this genetically regulated chain of events preceding death has any adaptive role or is just a suboptimal setting of stress-response machinery is unclear.

Here, we proposed that by undergoing cell death, yeast cells can protect their neighbouring cells against some naturally occurring xenobiotics. To test this hypothesis, we screened the effects of a number of xenobiotics and environmental stressors on prototrophic cells while supplementing them with viable or inviable auxotrophic yeast cells. We found that dead yeast cells inhibited the cytotoxic action of macrolide antifungal AmB. Furthermore, supplementation of yeast suspension with an AmB-sensitive strain can increase the average survival of cells in this suspension upon exposure to a high concentration of AmB. Together, our data show that, under certain conditions, decreased xenobiotic resistance in a subpopulation of cells can be beneficial for the microbial community.

## Results

Some stressors are better tolerated by yeast cells in dense communities than yeast cells less densely dispersed, whereas other stressors kill cells irrespective of the cell suspension density. To distinguish which stressors fall into which category, we subjected prototrophic (*HIS+*) yeast cells (5 × 10^6^ cell/ml) to various stresses in the absence or presence of histidine auxotrophs (*his*−). After stress was induced, cell mixtures were transferred to selective yeast nitrogen base (YNB) plates without histidine, where only *HIS+* cells can grow into a colony. We supplemented *HIS+* cells with either living histidine auxotrophs (live aux cells) or inviable histidine auxotrophs (dead aux cells, Figure 1). We performed heat shock to kill the auxotroph cells, because this method does not require additional procedures (e.g. centrifugation) to remove the stressor. Moreover, to assess cell suspension density effects, we tested equal and nine-fold higher cell density of histidine auxotrophs. We selected the concentration of xenobiotics/intensity of environmental stressors so that they kill at least 50% of *HIS+* cells under the control conditions, i.e. without the addition of auxotrophic cells.

**Figure 1.**
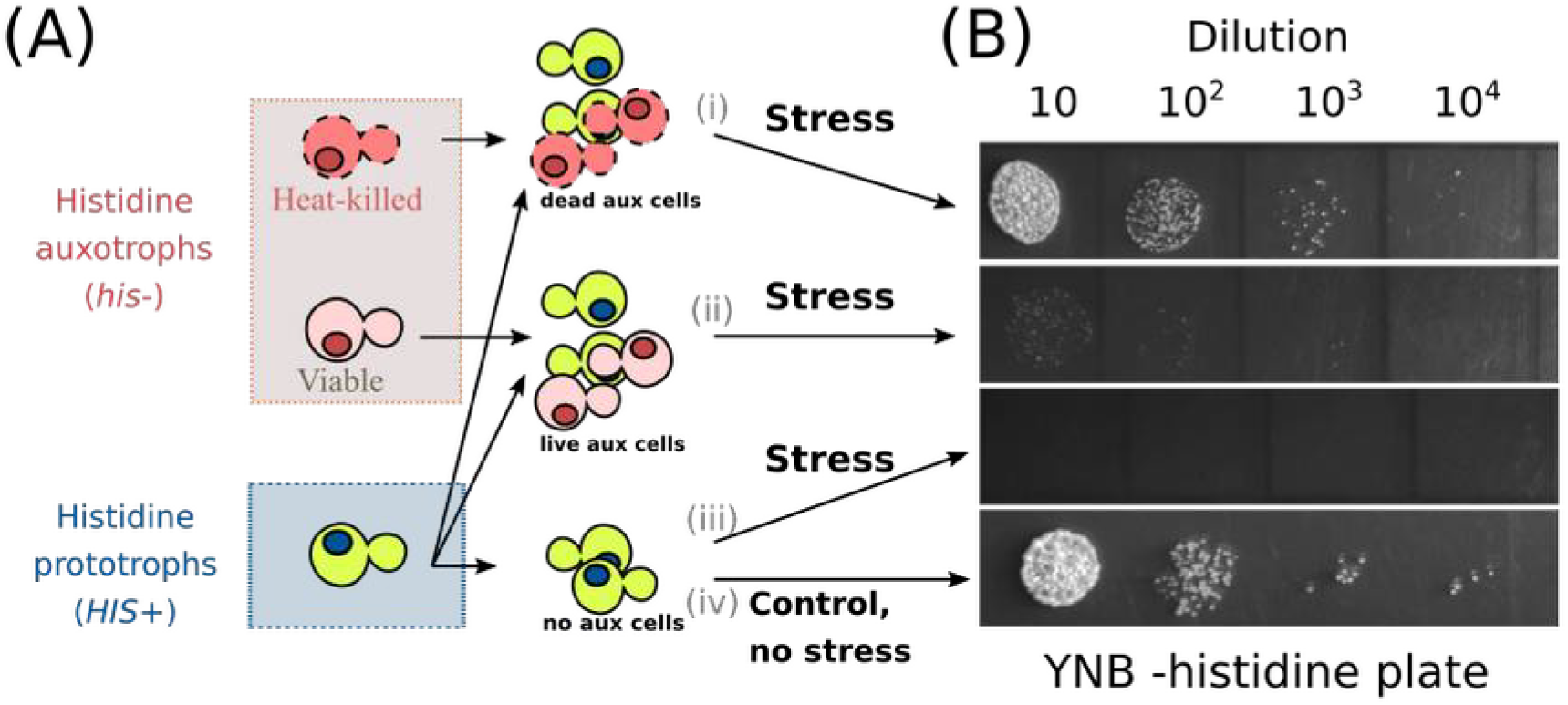
Scheme of experiment to test how an excess of viable and inviable auxotrophic cells alter the survival of prototrophic cells under various stress conditions. (A) Mixtures of (i) histidine auxotrophic (*his*−) heat shock-killed cells and histidine prototrophic (*HIS+*) living cells; (ii) histidine auxotrophic (*his*−) control cells and histidine prototrophic (*HIS+*) living cells; (iii) histidine prototrophic living cells were subjected to various stressors and transferred to an YNB-histidine medium plate. Unstressed control cells (iv) were plated before supplementation of stress factors. (B) Representative experiments in which AmB (7 μg/ml) was used as the stressor. In a typical experiment, we assessed 4.5 × 10^7^ cells/ml auxotrophic cells (*his*−) and 5 × 10^6^ cells/ml prototroph cells (*HIS+*).

Supplementation of viable or dead auxotrophic cells in most cases either increased the survival of *HIS+* cells or had no effect (Figure 2A). We classified stressors depending on how strong the protective effect of dead vs living cells was by clustering the responses of four tested conditions. The analysed types of stressors were clustered into two groups depending on whether the added auxotrophic cells helped the prototrophic cells to survive or not (Figure 2A). Strikingly, supplementation of dead auxotrophic cells protected prototroph yeast cells against macrolide antifungal AmB much better than an equal concentration of living cells. Stressors were ordered according to the relative effectiveness of the protection achieved by dead or living cells, which revealed that dead auxotrophic cells were the most effective in protecting prototroph from each of the tested AmB concentrations (Figure 2B). At the same time, in our dataset, we found no correlation between the relative efficacy of protection offered by dead cells and the average intensity of the stress (Figure 2C, Figure S1). Therefore, dead cells specifically increased survival in the case of AmB-induced death rather than increasing stress resistance in general.

**Figure 2.**
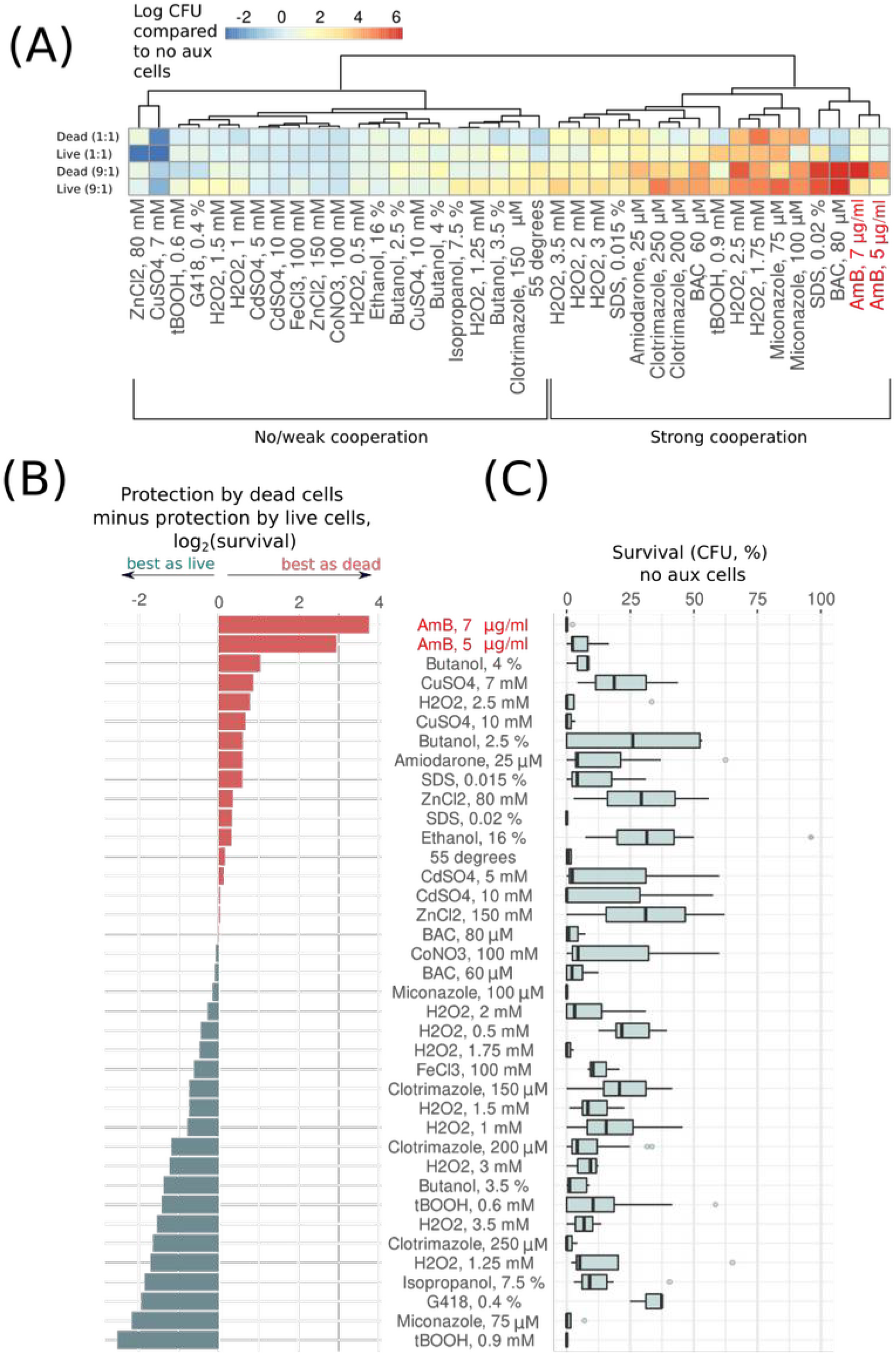
Excess of live and dead yeast cells in the suspension alleviates the lethality of some environmental stressors. (A) Heatmap indicates relative survival of prototrophic yeast cells supplemented with different amounts of living or heat-shock killed auxotrophic cells. (B) Stressors are ranked according to whether dead or living prototroph cells contribute more or less to the survival of the auxotroph’s suspension. Zero value indicates equal contribution of dead and living cells to the community stress resistance. (C) The panel shows the survival of prototroph yeast cells under the indicated stress conditions without the addition of auxotrophic cells (no aux cells).

To confirm the effect of dead cells on AmB-induced death in independent experiments, we subjected *his*− yeast cells to heat shock at different intensities. As a result, we obtained yeast suspensions of *his*− cells with different proportions of viable and inviable cells. We supplemented these *his*− yeast suspensions to *HIS+* cells and assessed the resistance of *HIS+* cells to AmB. In agreement with the result of our initial survey, the proportion of *his*− dead cells in the cell suspension correlated with the survival of *HIS+* cells treated with AmB (Figure 3A). Moreover, dead yeast cells protected yeast suspensions against other macrolide antifungals: filipin, natamycin and nystatin (Figure 3B).

**Figure 3.**
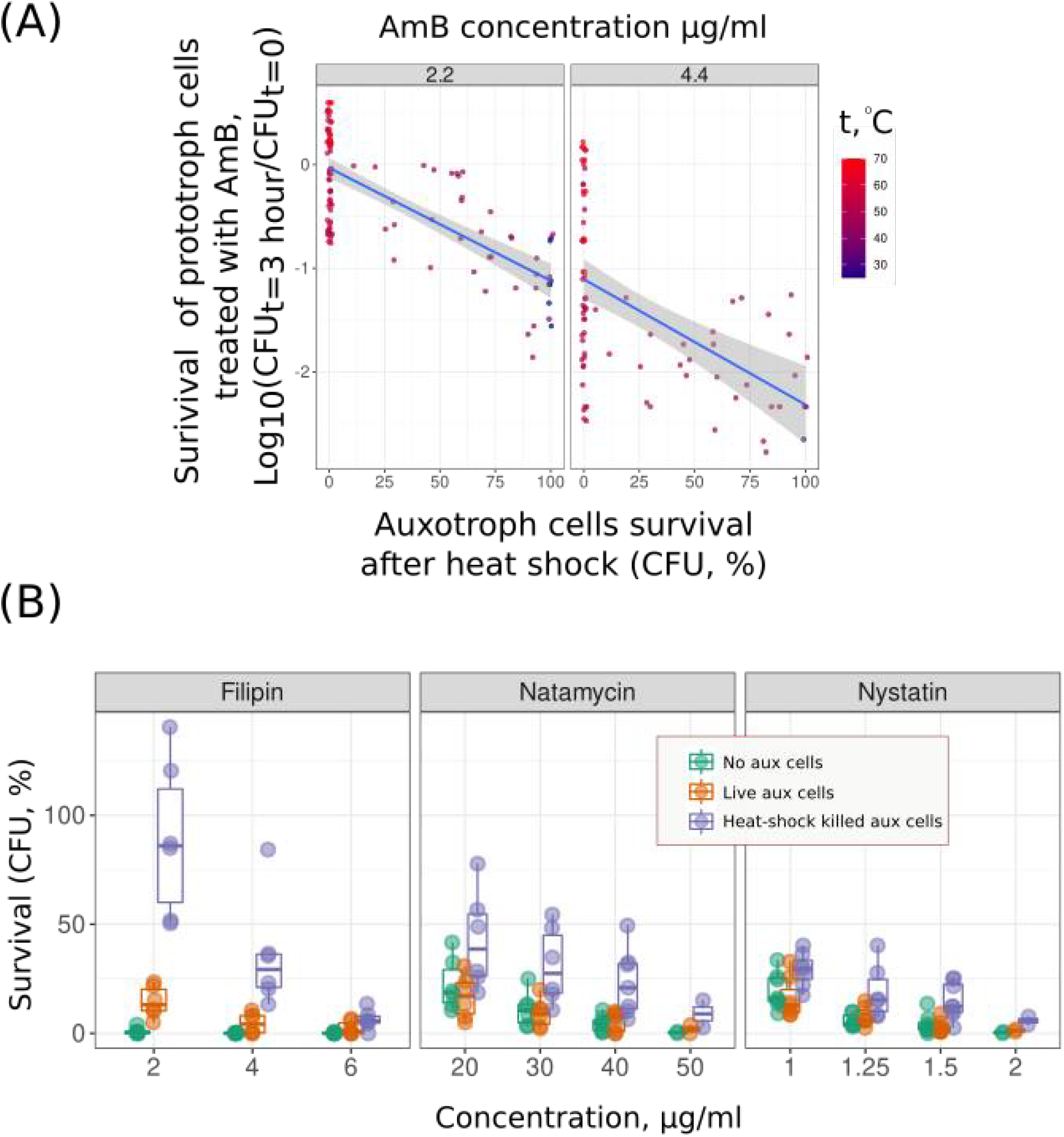
Supplementation of dead yeast cells protects yeast suspension against macrolide antifungals better than supplementation of additional living cells. (A) Protection of prototrophic yeast cells against AmB by auxotrophic cells killed by heat shock of different intensities. The protection provided by heat-shocked auxotrophic yeast cells to prototrophic yeast cells is proportional to the percentage of inviable auxotrophic yeast cells in the suspension. (2.2 μg/ml AmB: Kendall’s tau = 0.54, p-value = 1.77 × 10^−13^; 4.4 μg/ml AmB: tau = 0.443 p-value = 1.76 × 10^−7^); To perform this experiment, we treated auxotrophic (*his*− or *trp*−) yeast cells with different temperatures (30°C–70°C), added them to the corresponding prototrophic (*HIS+* or *TRP+*) strain, and then subjected them to AmB for three hours. (B) Supplementation of heat-shock killed cells increased the survival of *HIS+* prototrophic cells treated with macrolides: filipin, natamycin and nystatin.

Given that dead cells showed higher efficiency in protecting remaining surviving cells against macrolides, we reasoned that the supplementation of yeast suspensions with AmB-hypersensitive cells could increase the proportion of surviving cells. To test this hypothesis, we tested *Δpmp3* and *Δlam1Δlam2Δlam3Δlam4* strains that were previously shown to be sensitive to AmB (26, 27). We confirmed that AmB inhibited the growth of these strains at low concentrations, which did not prevent the growth of the parental strains (Figure 4A). Then, we treated yeast suspensions composed of the wild-type histidine prototrophic strain (*HIS+*, cell density 2 × 10^7^ cells/ml) and either of these strains. As a control, we supplemented the wild-type histidine prototroph with the parental auxotrophic strain (*his*−, see the scheme of experimental design in Figure 4B). We additionally supplemented the *Δlam1Δlam2Δlam3Δlam4 his*- cells but not the parental *his*- control cells that protected the wild-type yeast *HIS+* strain from a high concentration of AmB (Figure 4C) and filipin (Figure S2). The protection effect increased with the increase in auxotroph cells density. In the case of *Δlam1Δlam2Δlam3Δlam4 his-* cells, the protection of *HIS+* was already pronounced at a cell density 4 × 10^7^. *Δpmp3 his-* cells increased AmB-resistance of *HIS+* cells when added at a concentration nine times that of *HIS+* cells (Figure 4C, OD = 18, equivalent of 3.6 × 10^8^ cells) but only marginally.

**Figure 4.**
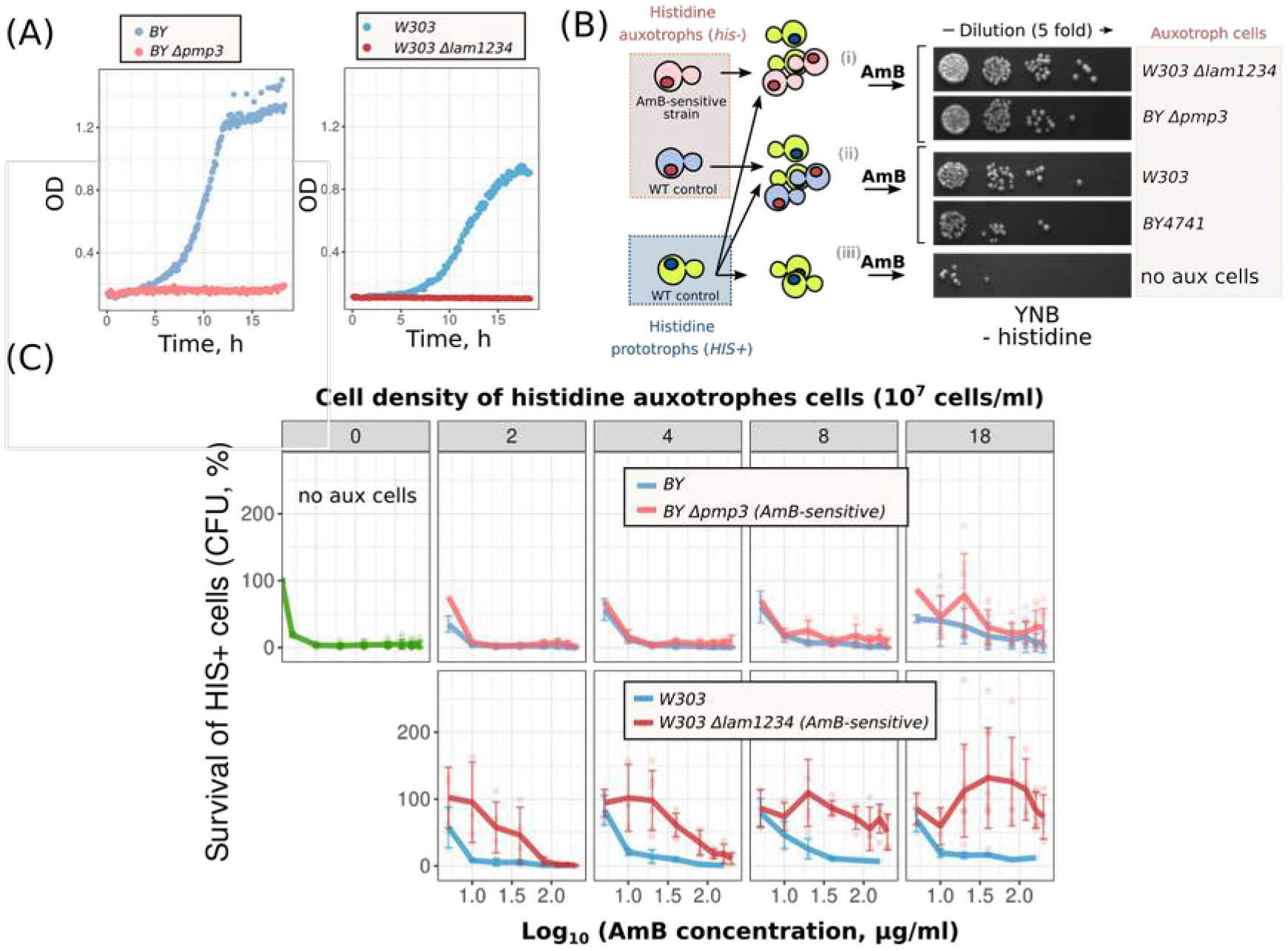
AmB-sensitive cells *Δlam1Δlam2Δlam3Δlam4* protect wild-type yeast cells from AmB better than the same amount of control cells. (A) Growth of *Δpmp3*, *Δlam1Δlam2Δlam3Δlam4* (*lam1234*) and control cells in the presence of AmB (0.8 μg/ml). (В) Scheme of the experiment. (C) Survival of *WT HIS+* cells treated with AmB. *WT HIS+* cells 2 × 10^7^ cells/ml were supplemented with auxotrophic strain either *Δpmp3 his-* or control *BY4741 his-* cells (upper panel); either *Δlam1Δlam2Δlam3Δlam4* (*lam1234*) or *W303* control (lower panel). Concentrations of auxotrophic cells are indicated in the upper row of the panel.

Strikingly, we found that the suspension when consisting of equal proportions of *Δlam1Δlam2Δlam3Δlam4* cells (2 × 10^7^ cells/ml) and control cells (2 × 10^7^ cells/ml) produced more colony-forming units (CFUs) on rich medium than the control cell suspension (4 × 10^7^ cells/ml) if treated with the same concentration of AmB (Figure 5). This result means that each AmB-sensitive cell that was killed by AmB saved more than one control cell with the wild-type AmB resistance.

**Figure 5.**
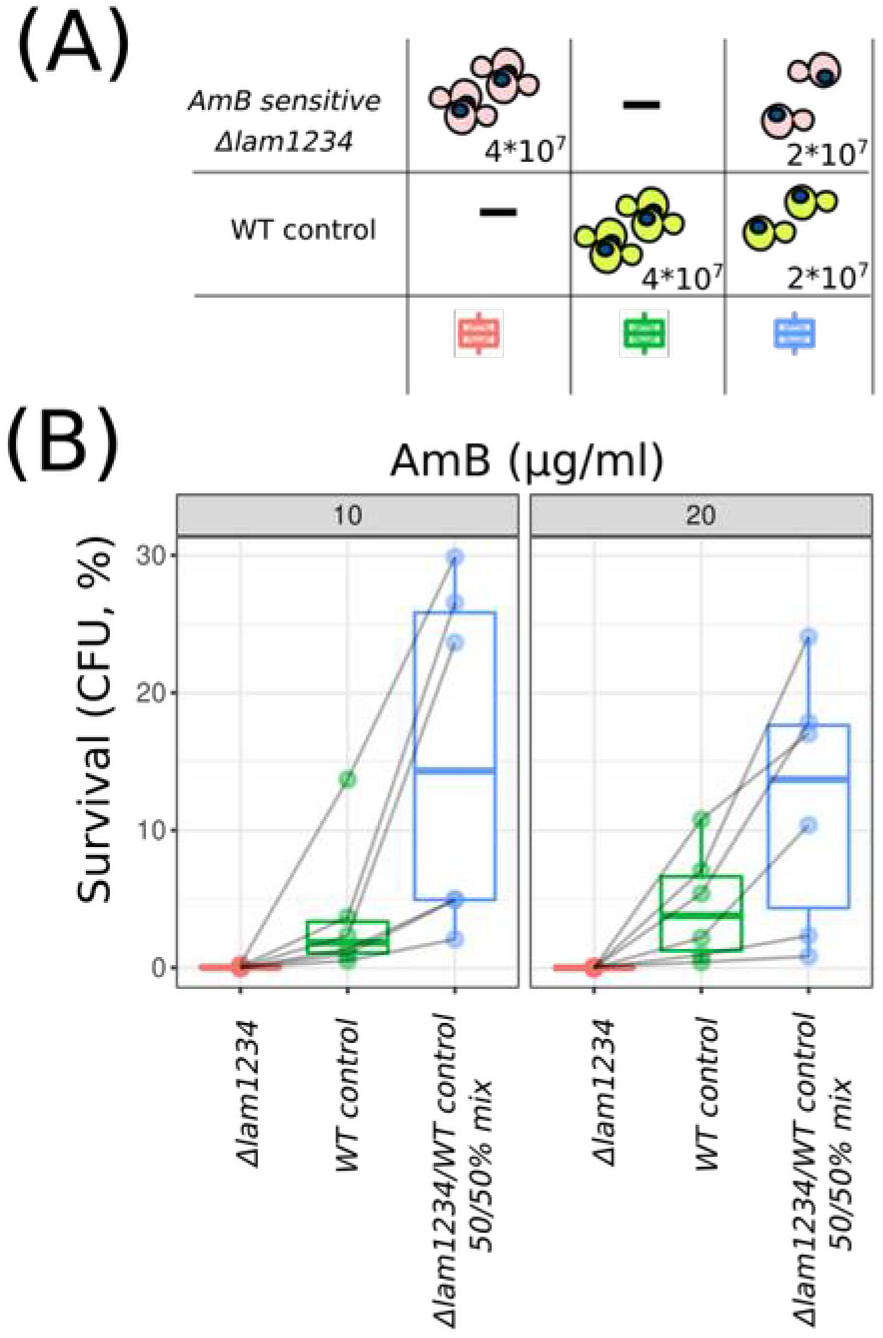
Cell mixture of WT and AmB-sensitive cells survive AmB better than homogenic WT cells. (A) Scheme of the experiment and figure legend. In all cases, we equalised the final concentration of cells in the testing tubes. Numbers designate the final concentration of cells in ml. (B) Average yeast cell survival in the wild-type (W303), Δlam1234 and the wild-type/Δlam1234 mixed suspensions treated with 10 or 20 μg/ml of AmB. In these experiments, we assessed cell survival by calculating the number of CFUs in YPD plates after three hours of AmB treatment (e.g. we did not distinguish the strain of surviving cells from the mixed suspensions). Shaded gray lines connect data points from separate day experiments. *P* = 0.027 according to paired Wilcoxon signed rank test for a comparison of the wild-type/Δlam1234 mixed suspension with the wild-type suspension.

We tested two possibilities to obtain insight into the mechanisms by which dead cells can protect living cells. First, it was shown earlier that AmB toxicity is mediated by secondary oxidative stress (28) and can be alleviated by the supplementation of antioxidants (29). We suggested that dead cells can release catalase from their cytoplasm into the incubation media and in this way protect living cells from AmB. To test this possibility, we put a genomic copy of cytoplasmic catalase gene *CTT1* under the regulation of a P_GAL_ promoter (see Materials and Methods) and overexpressed it by growing yeast in a galactose-containing medium. P_GAL_-*CTT1 trp-* catalase overexpression cells showed increased resistance to hydrogen peroxide (Figure 6A) but provided no increase in survival to corresponding prototrophic *TRP1+* cells subjected to amphotericin (Figure 6B).

**Figure 6.**
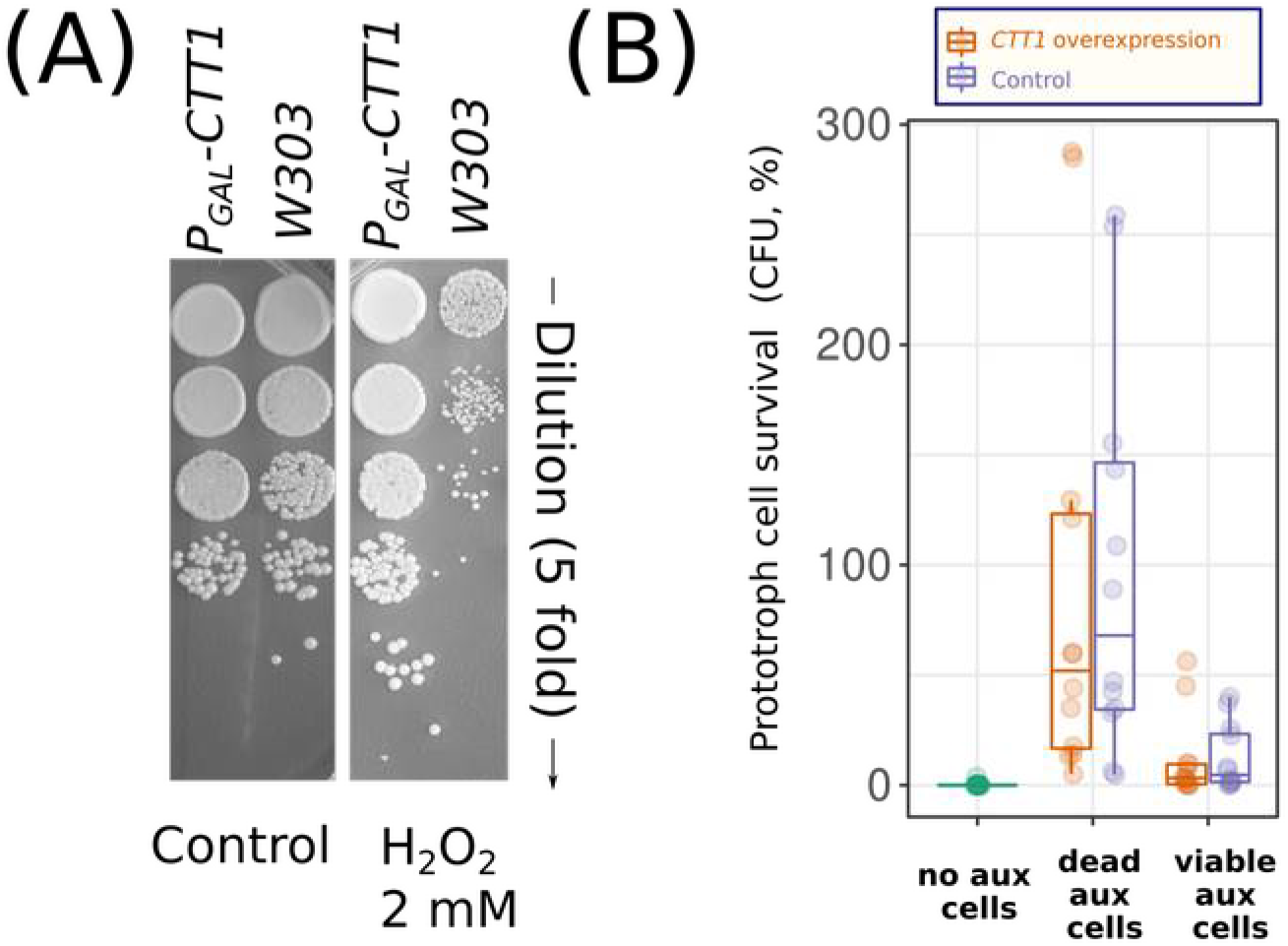
Overexpression of *CTT1* in tryptophan auxotrophic cells provides no increase in the survival of *TRP1+* cells in the same suspension. (A) Overexpression of *CTT1* provides resistance to hydrogen peroxide (3 h incubation times). To increase *CTT1* expression, we incubated the P_GAL_-*CTT1* strain in galactose-containing rich medium (YPGal) overnight. (B) Experimental design as described in Figure 1 with the only difference being that we used tryptophan selection instead of histidine. Cells were treated with 7 μg/ml AmB; the incubation time with AmB was three hours.

The second possibility we considered was that dead yeast cells absorb macrolide from the medium and, therefore, decrease the amount of antifungals bound to the membranes of living cells. Given that filipin has a high fluorescent yield and binds to sterol-rich membranes, it is a popular reagent to use for visualising sterol-rich membranes in yeast cells (27, 30). To test whether inviable yeast cells become permeabilised and absorb more macrolide, we stained the control and heat-shocked killed cells with filipin. Figure 7A shows that while control cells were stained only at the periphery, the heat-shocked cells exhibited intracellular compartment staining. To quantify the absorption of filipin by dead vs living cells, we incubated the yeast suspensions with filipin (5 μg/ml), centrifuged the suspension and measured the residual fluorescence in the supernatant (see Materials and methods section for details). The addition of dead cells to a filipin-containing incubation medium decreased the fluorescence in the supernatant (Figure 7B). Suspension of the dead yeast cells (OD = 5, equal to 10^8^ cells/ml) absorbed at an average filipin concentration of 2.55 μg/ml. At the same time, the same concentration of living control cells decreased the concentration of filipin in the supernatant by only 0.42 μg/ml.

**Figure 7.**
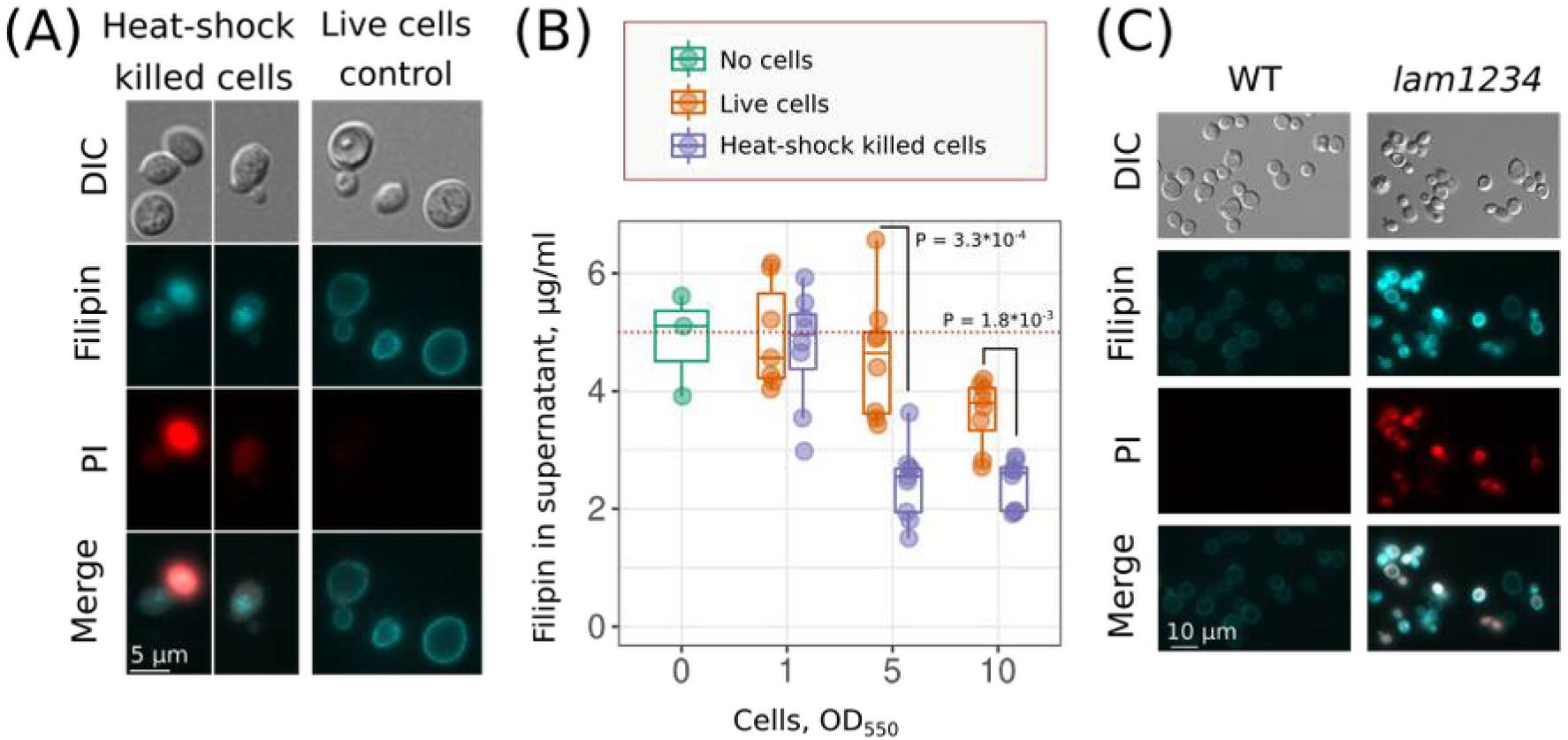
Dead yeast cells absorb macrolide filipin with intracellular compartments. (A) Different localization of the filipin signal in heat-shocked killed and live control cells; (B) Heat-shock killed cells absorb more filipin compared to viable control cells. Suspension of yeast cells was supplemented with filipin (5 μg/ml), then centrifuged. Integral fluorescence spectra in the supernatant were measured. *P* values were calculated according to the unpaired Mann–Whitney test. (C) Filipin staining induced permeabilization of *Δlam1Δlam2Δlam3Δlam4* (*lam1234*) strain but not the wild-type strain. Yeast cells were treated with filipin (5 μg/ml, incubation time 5 minutes).

Next, we treated the wild-type cells and *Δlam1Δlam2Δlam3Δlam4* mutant cells with filipin. In these cells, we analysed filipin intracellular localization. To test the integrity of the plasma membrane, we used propidium iodide, which is commonly used to detect dead cells. Figure 7C shows that five minutes of treatment of yeasts with filipin induced propidium iodide staining and intracellular filipin localization in *Δlam1Δlam2Δlam3Δlam4* cells but not in the control *W303* cells.

## Discussion

Cooperation among neighbouring cells can increase their resistance against some stressors but can be futile against the others. In the case of xenobiotics, hydrophobicity of the molecule is one of the basic factors determining the efficiency of cellular cooperation against it. Indeed, xenobiotics accumulating in cell membranes and lipid droplets can be depleted from media if there are excess cells and limited sources of xenobiotics. Accordingly, our survey of stressors (Figure 2A) showed that supplementation of additional auxotroph cells increased the survival of prototroph cells towards hydrophobic azole antifungals (e.g. miconazole) and surfactants (e.g. BAC) but did not alter their survival in the presence of heavy metals (e.g. CdSO_4_) or fusel alcohols (e.g. butanol). Additional yeast cells in suspensions also increased resistance to high concentrations of hydrogen peroxide (Figure 2A), which was probably due to the contribution of cellular antioxidant systems to the decomposition of hydrogen peroxide.

Meanwhile, we found that only non-viable cells provide significant protection against macrolide antifungals AmB, filipin and, to a lesser extent, natamycin and nystatin (Figures 2B, 3C). All of these compounds are produced by different species of *Streptomyces*—a widespread gram-positive soil filamentous bacteria (31). The biosynthesis of some of these compounds is species-specific: nystatin (*S. noursei*), AmB (*S. nodosus*) and others that are less specific, such as natamycin (32). Macrolide antifungals bind sterol rich membranes, induce their conductance and/or deplete membrane sterol from the membrane while disturbing its vital properties (33, 34). The amphiphilic nature of the macrolides suggests that they cannot passively diffuse across the membrane; moreover, cell walls additionally restrain macrolide AmB absorption by yeast cells (35). Therefore, in the suspension of live cells, macrolides interact primarily with the outer leaflet of plasma membranes. Although the plasma membrane contains more sterol than the membranes of other organelles (36), its surface area is much smaller than the integral surface of the cell membranes of permeabilised cells.

Moreover, some studies suggest that sterols are unevenly distributed within plasma membranes, with a major amount of sterol being available only from the inner (cytosol) leaflet of cytosol (37). For example, in yeast, only 20% of fluorescent dehydroergosterol (DHE) can be quenched by impermeable fluorescent quenchers, as efficient quenching requires the disruption of the plasma membrane integrity (38). Intriguingly, in our recent study, we found that the deletion of sterol-transporting LAM genes increase filipin staining of yeast cells in both the plasma membrane and intracellular compartments (27). The high sensitivity of the LAM-deficient strain to filipin (Figure 7C) suggests that intense plasma membrane staining in these experiments can be explained by filipin binding to the inner leaflet of yeast PM rather than an increase in the sterol concentration. Therefore, permeabilization of yeast cells can expose multiple additional macrolide antifungal-binding sites.

Given that permeabilised yeast cells absorb more macrolide antifungals than living yeast cells, a yeast community (e.g. dense suspension or colony) can benefit from early permeabilization of plasma membranes. This occurs in striking contrast to the apoptosis of mammalian cells, which maintains plasma membrane integrity to prevent the release of proinflammatory factors (39). Meanwhile, the metazoa in some cases rely on inflammation upregulation. Accordingly, during pyroptotic cell death of mammalian cells, plasma membrane rupture is facilitated by small plasma membrane proteins gasdermin D (40) and NINJ1 (41). Therefore, we speculate that the physiological scenarios of programmed cell death in yeast should be either homologous or analogous to metazoa programmed cell death mechanisms during early plasma membrane rupture.

Whether clonal microbial populations are heterogeneous is determined by the individual cells’ stress resistance phenotype (42, 43). This cell-to-cell heterogeneity arises from transcriptional noise, cell cycle-mediated differences and, in the case of budding yeasts, division asymmetry (44, 45). An increase in the variance of stress-resistance-phenotypes among individual cells in the population can improve the survival of clonal lineages through repetitive severe stresses (46). Our data extends these observations by exemplifying that improved survival in a suspension can be achieved by an increase in the variance of macrolide tolerance, even if this increase is associated with a decrease in the average tolerance. Indeed, Figures 4 & 5 show that the substitution of the control cells in a suspension with AmB-sensitive cells increases overall survival. We suggest that macrolide-resistance heterogeneity can be an adaptive trait that evolves to help cellular clonal communities to withstand a high concentration of macrolides.

## Materials and Methods

### Yeast strains, growth medium and reagents

We used standard yeast-rich and synthetic mediums described by Sherman (47). Yeast strains used in the study are listed in Table S1. To generate a strain with *CTT1* overexpression, we substituted the native *CTT1* promoter with a gene cassette containing the P_GAL_ promoter and a marker gene. To produce the cassette, we used the polymerase chain reaction (PCR)-based approach described by Longtine et al. (1998) (48) using *pFA6aHIS3MX6-PGAL1* plasmid as matrix DNA and the following gene-specific primers: *CTT1-F* 5′-ctcaatcttgtcgttacttgcccttattaaaaaaatccttctcttgtctcgaattcgagctcgtttaaac-3′, *CTT1-R* 5′-tttttaccgaacacgttcatttgtgaagctgagctgattgatcttattggcattttgagatccgggtttt-3′. Primers *CTT1-test-F* 5′-aatgatgagtacgtgcccgat-3′ and *CTT1-test-R* 5′-caccttcaagaggtttaggaa-3′ were used to validate the correct clones due to the size of the PCR product.

### Experimental setup for studying the contribution of heat-shock killed auxotroph cells to the survival of prototroph living cells

The cells were incubated overnight in 50-ml tubes with 5 ml liquid synthetic medium with all amino acids (YNB complete) at a cell density of 4 to 8 × 10^6^ cells/ml (logarithmic growth stage). The cells were collected by centrifugation (700 g, 5 min), and the medium was replaced with a synthetic medium without histidine (YNB–his). The experiment was performed in a 96-well plate with a cell mixture volume of 200 μl per well. We added to each well the cells of prototrophic and auxotrophic strains in one of several combinations: (1) 4 × 10^6^ cells/ml *HIS+* control cell; (2) 4 × 10^6^ *HIS+* cells/ml mixed with either live or heat-shock killed cells at 4 × 10^6^ cells/ml *his*− cells (prototrophic: live *his*− = 1:1, prototrophic: dead *his*− = 1:1); (3) 4 × 10^6^ cells/ml *HIS+* cells mixed with either live or heat-shock killed cells at 3.6 × 10^7^ cells/ml *his*− cells (prototrophic: live *his*− = 1:9, prototrophic: dead *his*− = 1:9). Then, xenobiotics or other stress factors were added, and the plate was incubated at 30°C and 700 rpm for three hours. Each sample was diluted 75 times and suspended; then, 5 μl of the diluted suspension was transferred onto plates with YNB-his. CFUs were counted in 24–48 hours (see the scheme of the experiment in Figure 1A and experimental results in Figure 2A). Figure 2 includes some experiments performed with four-fold lower cell density. Moreover, in some of the experiments shown in Figure 2, we used the *TRP+*/trp− prototroph/auxotroph pair.

We did not consider the possibility of histidine or tryptophan auxotrophy reversion in our experiments, given that no single colony had formed in the YNB-his and -trp plates when supplemented with the corresponding prototrophic strain cells.

To generate a set of auxotrophic cells with varying proportions of dead cells, the *W303* strain was exposed to different temperatures (30°C–70°C) for 30 minutes. Auxotrophic cell survival was defined as the ratio of CFUs after to before heat treatment. The set of auxotrophic cells (6 × 10^6^ cells/ml) with varying proportions of dead cells was mixed with exponentially growing prototrophic (*TRP+*) cells (1.2 × 10^6^ cells/ml). AmB at a concentration of 2.2 μg/ml or 4.4 μg/ml was added to the resulting mixtures; then, the mixtures were incubated for three hours at 30°C temperature and 200 rpm. The cells of the mixture were transferred onto plates with YNB-trp selective media. CFUs were determined after two days of growth at 30°C.

### Growth kinetics

Exponentially growing cells were diluted to an optical density of OD_550_ = 0.2 and inoculated into a 48-well plate (Greiner). Plates were incubated at 30°C in a spectrophotometer (SpectrostarNANO) with the following settings: orbital shaking at 500 rpm for 30 sec before measurements; OD_550_ measurements were performed at 5-min intervals.

### Fluorescent microscopy

We resuspended wild-type and mutant yeast cells in 50 mM potassium phosphate buffer to a final concentration 5 × 10^7^ cells/ml and supplemented the suspension with filipin (filipin complex from *Streptomyces filipinensis*; Sigma F9765) to a final concentration of 5 μg/ml. After five minutes of incubation, filipin was removed from the medium by centrifugation and cells were supplemented with propidium iodide (Thermo Fisher Scientific, P3566, final concentration 1 μg/ml). To photograph cells, we used an Olympus BX41 microscope with a U-MNU2 filter (excitation wavelength 360–370 nm; beam splitter filter 360–370 nm; emission > 420 nm) for filipin and U-MNG2 filter (excitation 530–550 nm, beam splitter filter 570 nm; emission > 590 nm) for propidium iodide. Photographs were taken with a DP30BW charged-coupled device camera.

### Filipin absorption experiments

Cells were grown overnight on solid YNB complete medium and then resuspended in 50 mM potassium phosphate buffer, pH 5.5, to a final OD_550_ of 1,5 or 10. Filipin was added to a final concentration of 5 μg/ml. After 5 min incubation, the cells centrifuged and supernatant was transferred to a 96-well plate. Fluorescence of unabsorbed filipin was analysed using Fluoroskan Ascent (excitation 355 nm; emission 460 nm). The filipin calibration curve, as shown in Figure S3, revealed the linearity of the tested concentration of filipin.

### Data analysis

We analysed data and generated the figures with R tidyverse libraries (49). A heatmap was generated with the pheatmap R package (version 1.0.12) with default parameters that use maximum linkage clustering. Where possible, we have shown individual data points and provided connections between data points obtained from the same experiment.

## Funding

The study was supported by the Russian Foundation for Basic Research grant N 18– 04-01183 (Figures 1–2) and the Russian Science Foundation grant N 18-14-00151 (Figures 3–7).

## Acknowledgements

We are grateful to Emir Radkevich, who participated in some of the preliminary experiments of this project. We are grateful to the G. F. Gause Institute of New Antibiotics, Moscow, Russian Federation, for providing us with nystatin. This research has been supported by the Interdisciplinary Scientific and Educational School of Moscow University Molecular Technologies of the Living Systems and Synthetic Biology.

## Notes

### Competing Interest Statement

The authors have declared no competing interest.

