## Supplementary materials for "The adaptive role of cell death in yeast communities stressed with macrolide antifungals"

Tables S1. Strains uses in the study

| **Strain** | **Genotype** | **Parental strains and/or references** |
|---|---|---|
| *W303-1A* | *MATa ade2-101 his3-11 trp1-1 ura3-52 can1-100 leu2-3* | Laboratory of A. Hyman |
| *HIS3* | *MATa ade2-101 his3-11 trp1-1 ura3-52 can1-100 leu2-3 HIS3* | Galkina et al. (2020) (50) |
| *TRP1* | *MATa ade2-101 his3-11 trp1-1 ura3-52 can1-100 leu2-3 TRP1* | Galkina et al. (2020) (50) |
| *Δlam1Δlam2 Δlam3Δlam4* | *MATa ade2-101 his3-11 trp1-1 ura3-52 can1-100 leu2-3 MATa ade2-101 his3-11 trp1-1 ura3-52 can1-100 leu2-3 Δlam3::kanMX4 Δlam2::TRP1 Δlam1::NAT Δlam4::loxP* | Sokolov et al. (2020) (27) |
| *BY4742* | *MATalpha his3Δ1 leu2Δ0 met15Δ0 ura3Δ0* | *EUROSCARF* |
| *BY Δpmp3* | *MATalpha his3Δ1 leu2Δ0 met15Δ0 ura3Δ0 Δpmp3::kanMX4* | *EUROSCARF* |
| $P_{GAL1}$-*CTT1* | *MATa ade2-101 his3-11 trp1-1 ura3-52 can1-100 leu2-3 PGAL1-CTT1::HIS3* | *W303-1A* |

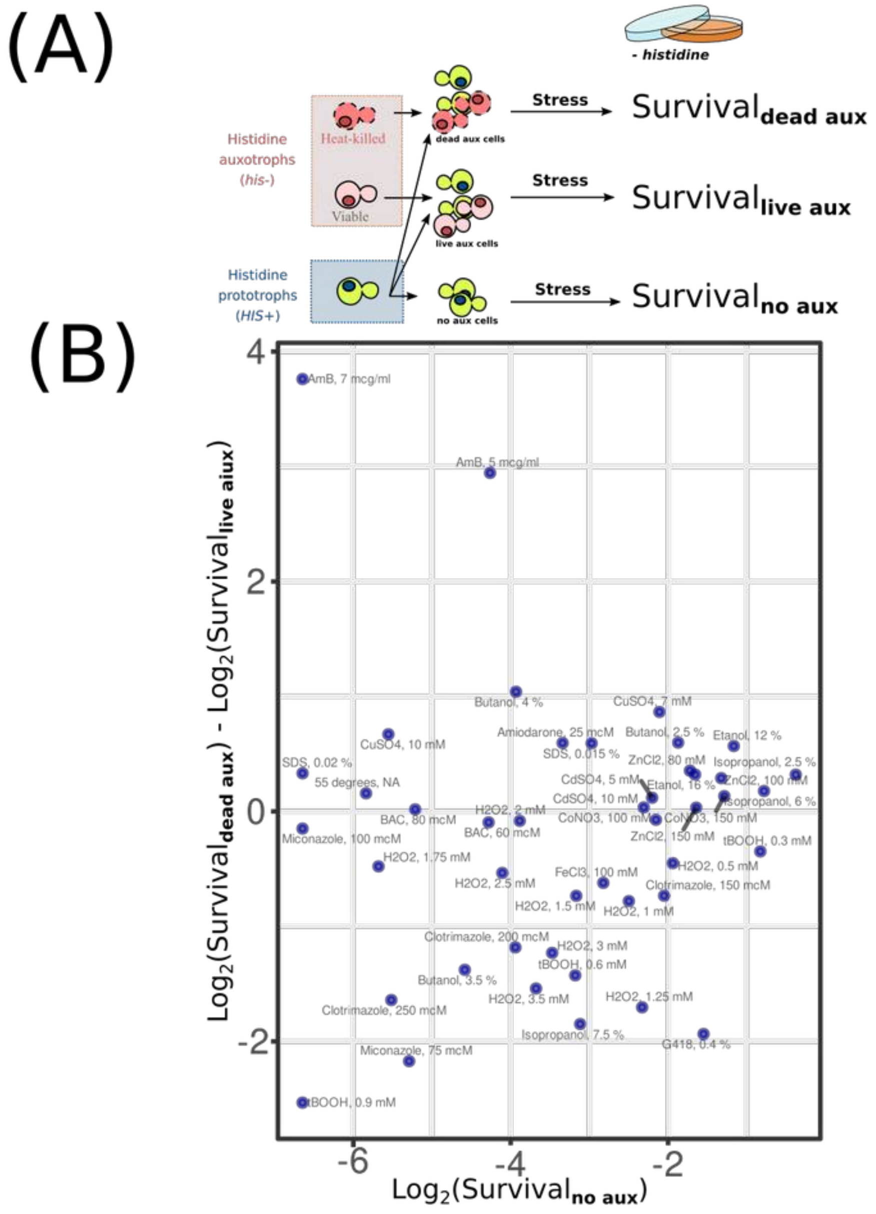


Figure S1. Stress intensity doesn't correlate with the relative effect of dead cells on survival of the prototrophic strain. (A) Scheme of the experiment (same as in figures 1,2); the survival of prototrophic cells was calculated as the number of CFU in suspension plated to YNB -histidine plate after the stress divided by the number CFU on YNB -histidine plated before the stress; (B) Y-axis: the relative survival of prototrophic strain supplemented with dead versul live auxotrophic cells ; X-cell: the survival of prototrophic cells without supplemental stress. The data is the same as in Figure 2B. Each data point corresponds to the average survival of yeast cells under specific stress. Kendall's tau = 0.1060451, z = 1.0119, p-value = 0.3116.

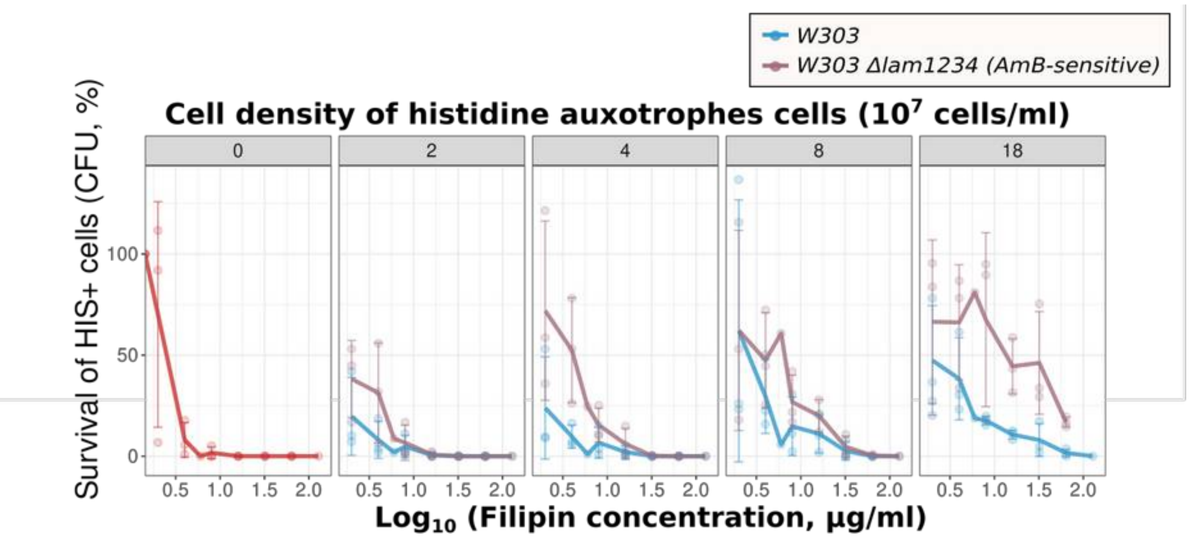


Figure S2. Survival of *WT HIS+* cells treated with AmB. *WT HIS+* cells $2*10^7$ cells/ml were supplemented with auxotrophic strain *Δlam1Δlam2Δlam3Δlam4* (*lam1234*) or *W303* control. Concentration of auxotrophic cells are indicated in the upper side of the panel.

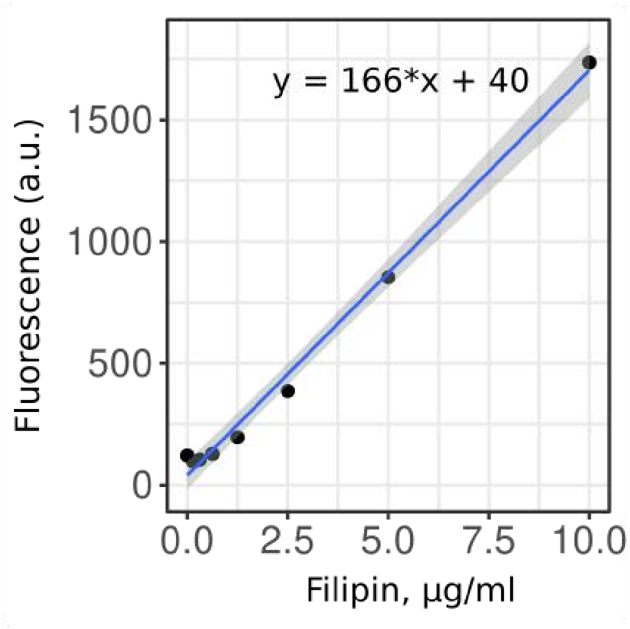


Figure S3. Calibration curve for filipin
